# Inhibiting ACSL1 related ferroptosis restrains MHV-A59 infection

**DOI:** 10.1101/2021.10.14.464337

**Authors:** Huawei Xia, Zeming Zhang, Fuping You

## Abstract

Murine hepatitis virus strain A59 (MHV-A59) belongs to the β -coronavirus and is considered as a representative model for studying coronavirus infection. MHV-A59 was shown to induce pyroptosis, apoptosis and necroptosis of infected cells, especially the murine macrophages. However, whether ferroptosis, a recently identified form of lytic cell death, was involved in the pathogenicity of MHV-A59, is unknown. Here, we demonstrate inhibiting ferroptosis suppresses MHV-A59 infection. MHV-A59 infection upregulates the expression of *Acsl1*, a novel ferroptosis inducer. MHV-A59 upregulates *Acsl1* expression depending on the NF-kB activation, which is TLR4-independent. Ferroptosis inhibitor inhibits viral propagation, inflammatory cytokines release and MHV-A59 infection induced cell syncytia formation. ACSL1 inhibitor Triacsin C suppresses MHV-A59 infection induced syncytia formation and viral propagation. *In vivo* administration of liproxstatin-1 ameliorates lung inflammation and tissue injuries caused by MHV-A59 infection. Collectively, these results indicate that ferroptosis inhibition protects hosts from MHV-A59 infection. Targeting ferroptosis may serves as a potential treatment approach for dealing with hyper-inflammation induced by coronavirus infection.

## Introduction

SARS-CoV-2, the causative agent of the coronavirus disease 2019 (COVID-19), has resulted in more than 4 million deaths worldwide. Thus, studying coronavirus and interaction between coronavirus and hosts can help understand the mechanism within the infection process and provide new approaches for clinical treatment. Murine hepatitis virus strain A59 (MHV-A59) is a well-studied coronavirus infection model (1). It has been validated that lung infection of MHV-A59 was sufficient to cause pneumonia and severe lung injuries (2, 3). Expression of inflammatory cytokines such as *Cxcl10, Ifng* and *Il6* were potently elevated in the lungs of infected mice. Mice with intranasal inoculation of MHV-A59 exhibited typical acute inflammatory response with large areas of hemorrhages. Our previous study also showed that intranasal MHV-A59 infection mouse model mimicked the hyper-inflammation induced by SARS-CoV-2 infection (4), further suggesting that intranasal inoculation with MHV-A59 can serve as a surrogate mouse model for studying COVID-19.

Ferroptosis is a newly identified iron-dependent necrotic cell death, mainly raised by the accumulation of lipid reactive oxygen species (ROS), resulting in excessive lipid peroxidation and subsequent cell membrane damage (5, 6, 7). Ferroptosis can be divided into two types, namely canonical and non-canonical types, according to their dependency on the involvement of GPX4. In the canonical ferroptosis, inactivation or depletion of GPX4 leads to the lack of glutathione (GSH) and lipid ROS overload, making the cell membrane vulnerable to lipid peroxidation (8). The non-canonical ferroptosis is characterized by the inactivation of ferroptosis suppressor protein 1 (FSP1), which maintains the active pool of radical-trapping antioxidant coenzyme Q10 (9, 10). In addition, cystine/glutamate transporter (xCT), encoded by *Slc7a11*, transfers cystine, the central material for generating GSH, into the cytosol to prevent ROS overload induced cell damage (11). Peroxidation of phospholipid species acts as the execution stage of ferroptosis. At the execution stage, for example, polyunsaturated fatty acids (PUFAs) can be esterified by acyl-CoA synthetase long-chain family member 4 (ACSL4), which occurs at the cell membrane, leading to the cell membrane rupture (12, 13, 14). Recently, acyl-CoA synthetase long-chain family member 1 (ACSL1) was uncovered as a ferroptosis promoter (15). ACSL1 promoted α-eleostearic acid (αESA) triggered ferroptosis. However, the αESA triggered ACSL1 dependent ferroptosis was distinct from the ferroptosis induced by the canonical ferroptosis inducers, including the GPX4 inhibitor ML160 and the FSP1 inhibitor iFSP1. ACSL1 has only been validated to participant in a murine breast cancer xenograft model (16). How ACSL1 involves in other biological processes is unknown.

Viral infection induces various kinds of programed cell death, including apoptosis, necroptosis and pyroptosis (17). For the β-coronavirus infection, cells were reported to undergo PANoptosis, which consisted of <u>P</u>yroptosis, <u>A</u>poptosis and <u>N</u>ecroptosis (18, 19). Previous studies have shown that SARS-CoV-2 can induce these three kinds of programed cell death in susceptible epithelial cells and immune cells (20, 21, 22). As for ferroptosis, whose morphological and biochemical features are distinct from PANoptosis, new castle virus (NDV) was the first virus to be discovered as ferroptosis inducer in tumor cells (23). NDV infection reduced the expression of *Slc7a11* and *Gpx4*, resulting in the ferroptosis of infected cells. Apart from NDV, so far, no other viruses were reported to induce ferroptosis of infected cells, although hepatitis B, hepatitis C, HIV-1 and human cytomegalovirus infection caused increased serum iron level (24, 25).

Ferroptosis has been linked with inflammation and immune responses. In innate immunity, GPX4 deficiency dampens cGAS-STING dependent innate immune signaling activation (26). Neutrophils from systemic lupus erythematosus (SLE) patients exhibit ferroptosis due to the transcriptional repression of *GPX4* (27). In adaptive immunity, GPX4 is essential for maintaining the metastasis of T_FH_ cells and promotes humoral immune responses (28). Ferroptosis of non-leukocytic cells often results in the release and activation of different damage-associated molecular pattern (DAMP), triggering the inflammatory responses (29, 30). Because COVID-19 patients suffered from systemic hyper-inflammation, characterized by ROS elevation and cytokine storm, SARS-CoV-2 infection was predicted to involve ferroptosis (31, 32, 33). The prediction was also evidenced by the elevated serum iron load in COVID-19 patients and decreased *GPX4* expression in SARS-CoV-2 infected cells (34, 35, 36). Here we showed that MHV-A59 infection induced *Acsl1* expression and ferroptosis of murine macrophages. Inhibition of ACSL1 with Triacsin C and ferroptosis inhibitors protected cells from MHV-A59 infection. Intranasal inoculation of liproxstatin-1 ameliorated MHV-A59 infection induced hemorrhagic alterations and immune cells infiltration in lungs of infected mice. Our study provides evidence for targeting ferroptosis to deal with coronavirus infection.

## Results

### Ferroptosis inhibitors suppressed syncytia formation after MHV-A59 infection

Coronavirus infection results in typical cell-cell fusion named syncytia (37). Syncytia formation of SARS-CoV-2 or MHV-A59 infected cells mainly depends on the interaction between spike protein and receptors and indicates the viral abundance and cellular antiviral mechanism (38). Although certain cell types such as lung epithelial cells and colorectal epithelial cells are susceptible to SARS-CoV-2, it has been reported that SARS-CoV-2 was able to infect macrophages (39, 40, 41). Both MHV-A59 and SARS-CoV-2 can infect murine macrophages. MHV-A59 infected murine bone marrow derived macrophages (BMDMs) and peritoneal macrophages (PMs) via the canonical receptor Ceacam1 (42), while Nrp1 mediated SARS-CoV-2 infection of murine macrophages and human olfactory epithelium (43, 44, 45). Given that coronavirus infection in macrophages promotes the inflammatory responses, which involves pro-inflammatory cell death programs, we intended to determine whether ferroptosis contributes to the pathogenicity of coronavirus. We first monitored cell morphology of both BMDM and PM after MHV-A59 infection for 24 hours. Cell-cell fusion resulted in multinucleated cell formation, followed by the appearance of squeezed out vacuole and subsequent cell death, which was ferroptosis-like (Video S1). It has been shown that syncytia were observed in tissues from deceased patients with pulmonary manifestations compared with those without pulmonary manifestations (38). We next used ferrostatin-1 (Fer-1), a ferroptosis inhibitor that functions via trapping radical, to check whether ferroptosis inhibition influenced MHV-A59 infection (46, 47). Syncytia formed by peritoneal macrophages was apparently inhibited by Fer-1 (Figure 1A and 1B). Besides, Fer-1 also protected cells from membrane damage (Figure 1C). In contrast, the caspase-3 inhibitor z-DEVD-FMK and caspase-1 inhibitor VX765 showed no obvious effect on cell membrane integrity maintenance, whereas the RIPK3 inhibitor GSK-872 exerted protective effects similar with Fer-1 (Figure 1D). Collectively, these results indicated that ferroptosis inhibition protected murine macrophages from MHV-A59 infection.

**Figure 1.**
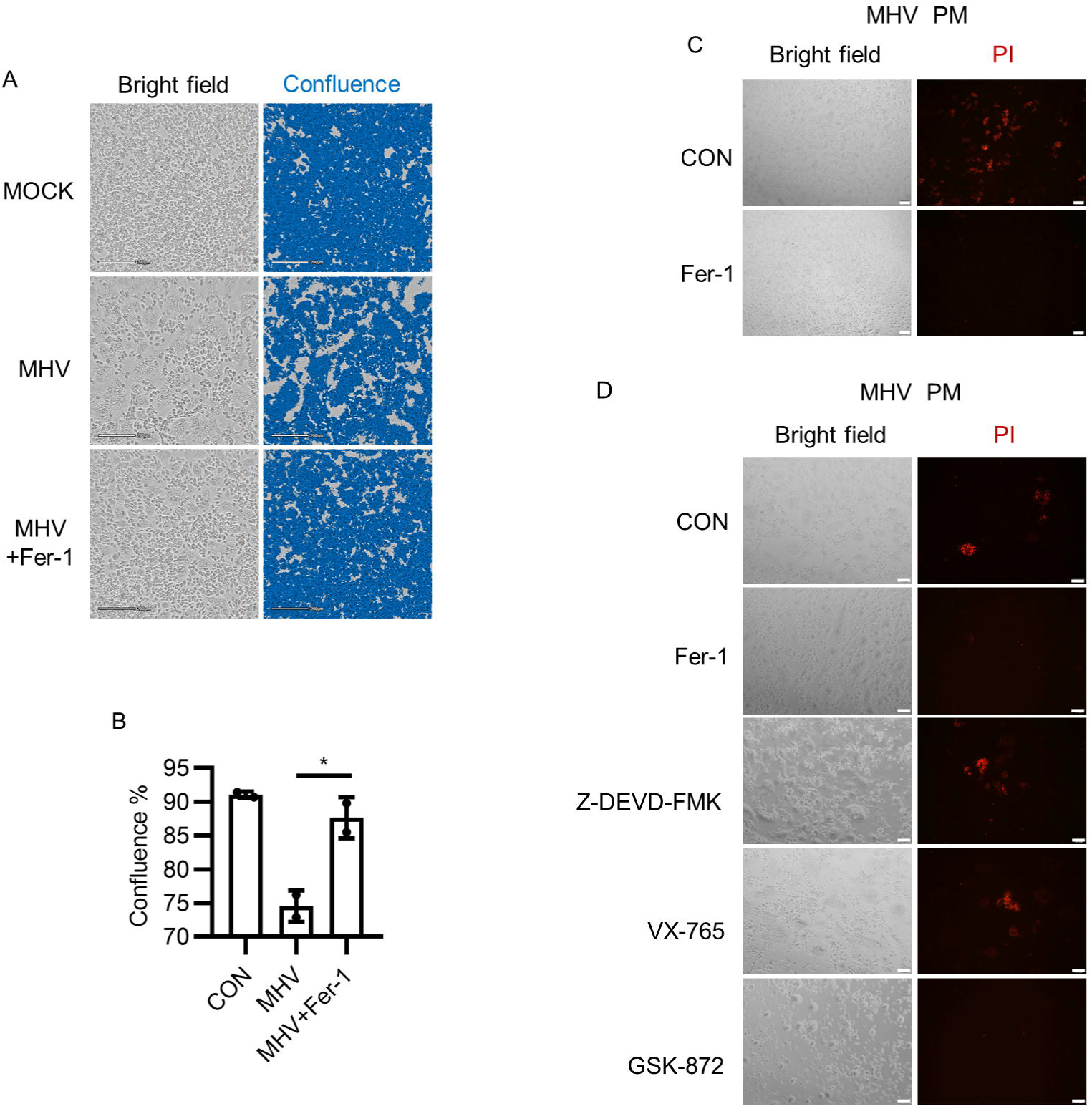
Ferroptosis inhibitor Fer-1 inhibited MHV-A59 induced syncytia and membrane damage. (A and B) Primary peritoneal macrophages (PMs) were infected with MHV-A59 at 0.05 MOI. After 2 hours of infection, cells were treated with Fer-1 (10 μM). After 24 hours of infection, cells were imaged using CytoSMART system (A) and cell confluence was evaluated using the CytoSMART website (B). (C) PMs were infected as described in (A) and were stained with propidium iodide (PI). After staining cells were imaged under fluorescence microscope. (D) PMs were infected as described in (A), followed by the treatment with Fer-1 (10 μM), z-DEVD-FMK (25 μM), VX-765 (30 μM) or GSK-872 (10 μM). Cells were stained with PI and imaged under fluorescence microscope. (E) Scale bars: For (A) and (C), 200 μm. For (D), 100 μm. Data from two independent experiments was shown. *, *p* < 0.05; Student’s *t-*test. See also Video S1.

### Ferroptosis inhibitors reduced viral load and inflammatory cytokine release after MHV-A59 infection

We next intended to find how ferroptosis inhibitors restrict MHV-A59 infection. We first tested genomic RNA level of MHV-A59 after 2 hours of infection. Fer-1 administration had minimal effects on invasion of MHV-A59 into macrophages (Figure S1). After excluding the possibility that Fer-1 inhibited viral entry, we next wanted to know whether Fer-1 inhibited MHV-A59 propagation. Interestingly, quantitative PCR results showed that intracellular MHV RNA level was identical or even slightly higher in cells treated with Fer-1 after infection (Figure 2A). However, via evaluating the viral titer in the cell culture supernatant, we found that viral load was lower after Fer-1 treatment (Figure 2B). These results suggested that ferroptosis inhibition affected viral propagation of MHV-A59. Because ferroptosis often occurs along with inflammatory cytokine release (48), we wanted to determine how ferroptosis inhibition influenced MHV-A59 induced inflammation. Fer-1 treatment showed no significant effect on the expression of inflammatory cytokines (*Il6*, and *Cxcl10*) and type I interferon (*Ifnb1*) (Figure 2C and 2D). However, Fer-1 treated macrophages released lower level of IL-6 (Figure 2E). These data suggested that ferroptosis inhibition reduced inflammatory cytokine release after MHV-A59 infection. LDH release level has been used to evaluate cell viability in lytic cell death. We thus also checked effects of Fer-1 on LDH release after MHV-A59 infection. As shown in Figure 2F, less LDH was released from Fer-1 treated cells, indicating the higher viability of cells after ferroptosis inhibition. Taken together, these data indicated that ferroptosis inhibition restricted propagation of MHV-A59 and inflammatory cytokine release induced by MHV-A59 infection.

**Figure 2.**
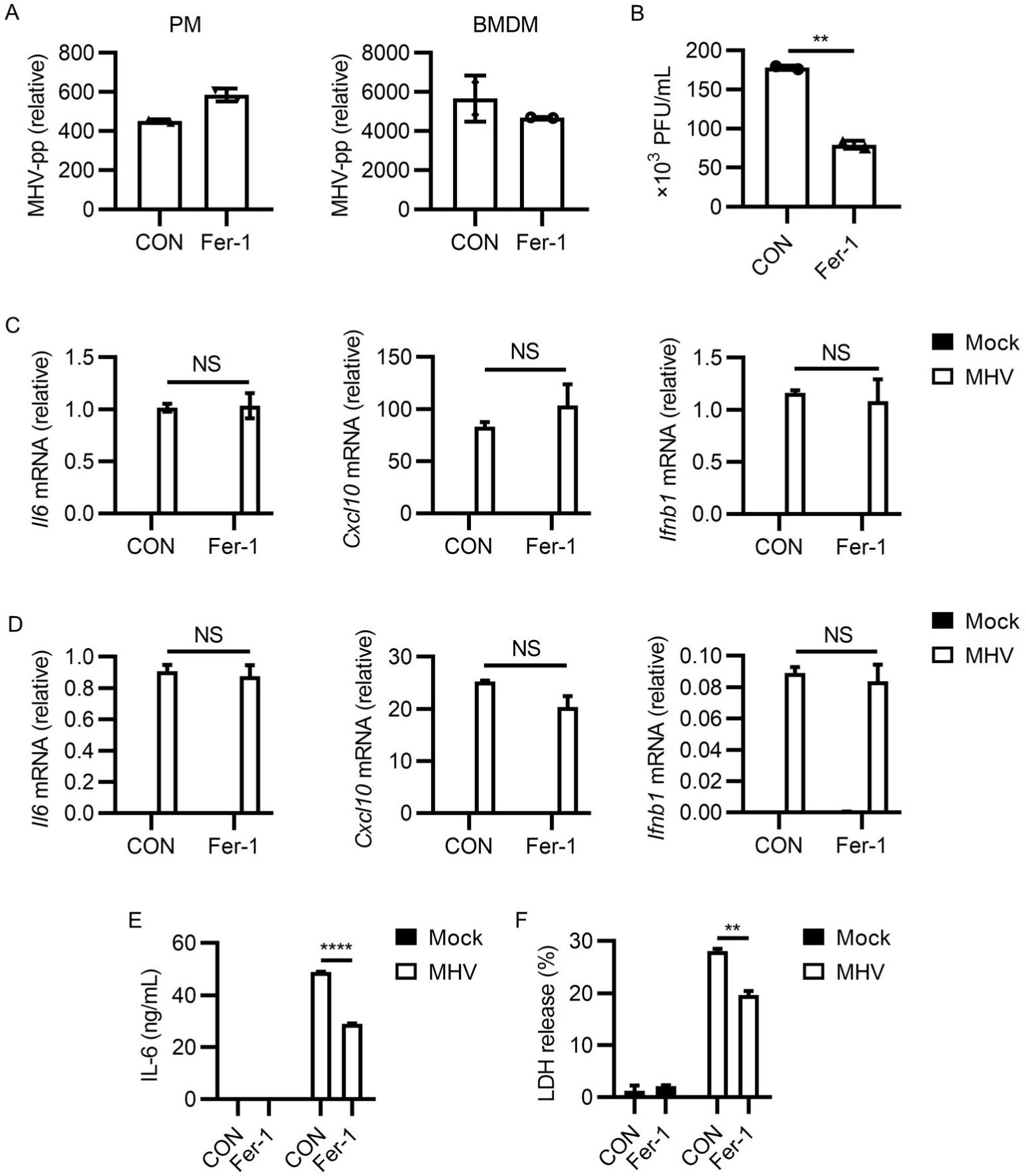
Fer-1 inhibited MHV-A59 propagation and infection induced inflammatory cytokine and LDH release. (A) PMs or BMDMs were infected with MHV-A59 at 0.05 MOI for 24 hours. Fer-1 was added after 2 hours of infection. MHV abundance was evaluated by the expression of MHV polyprotein (MHV-pp). MHV-pp expression was tested by qRT-PCR. (B) Viral load of MHV-A59 from supernatants of PMs from (A) was evaluated with plaque assay. (C and D) Expression of *Il6, Cxcl10* and *Ifnb1* after MHV infection in PMs (C) or BMDMs (D) with or without Fer-1 treatment was tested by qRT-PCR. (E) IL-6 abundance from supernatants of PMs from (C) was tested with ELISA. (F) Percentage of LDH release of PMs from (C) was tested by LDH detection kit. Data from two independent experiments was shown. **, *p* < 0.01; ****, *p* < 0.0001; Student’s t-test. NS, not significant. See also Figure S1.

### Liproxstatin-1 ameliorated MHV-A59 infection induced inflammation and lung injury

We next checked the efficiency of ferroptosis inhibitors on inhibiting MHV-A59 lung infection. We selected C57BL/6 mice to carry out the viral infection experiments, because intranasal inoculation of MHV-A59 was reported to cause severe lung inflammation and tissue injury in this mouse strain. Liproxstatin-1 (Lip-1) functions as another radical-trapping ferroptosis inhibitor and has been selected as ferroptosis inhibitor in several mouse models (8, 27). We thus assessed the influences of Lip-1 treatment on MHV-A59 lung infection. Daily intranasal treatment of Lip-1 was carried out after MHV-A59 infection until day 10 post infection (Figure 3A). Comparison of pathologic alterations in lungs of mice between MHV-A59 infection alone group and infection together with Lip-1 treatment group showed that Lip-1 treated group exhibited less severe pneumonia, characterized by less hemorrhagic alterations and less immune cells infiltration (Figure 3B, 3C and 3D). However, such model applying low dose of MHV-A59 resulted in no significant difference in body weight changes between groups with or without Lip-1 treatment after MHV-A59 inoculation (Figure S2). Taken together, these data suggested that Lip-1 ameliorated inflammatory responses and tissue injury in lung after MHV-A59 infection.

**Figure 3.**
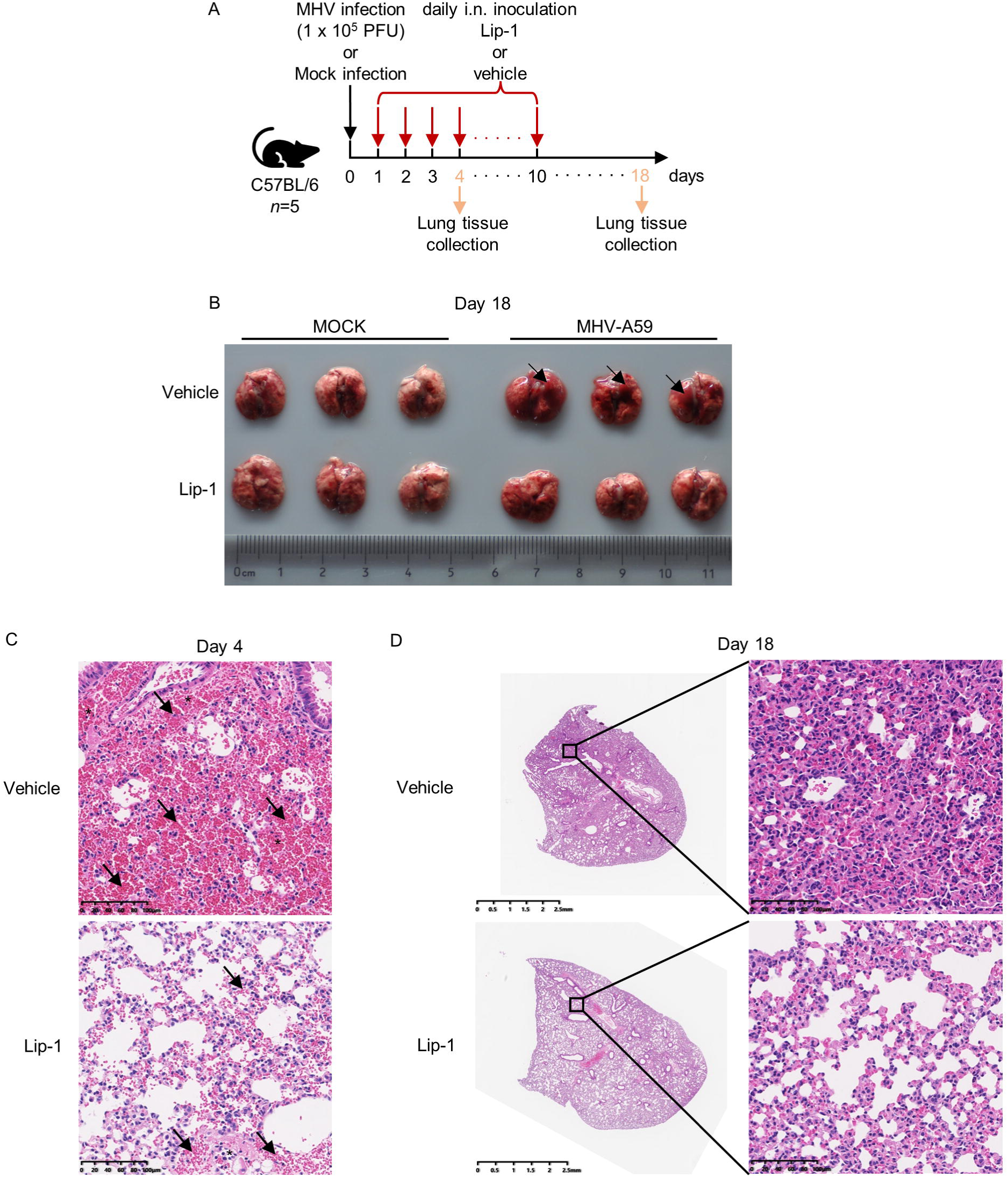
Lip-1 alleviated MHV induced lung tissue injury. (A) Diagram of MHV-A59 infection model. 4-week-old male C57BL/6 mice were infected intranasally (i.n.) with 1 × 10^5^ PFU MHV-A59 in 10 μL DMEM. Lip-1 (10 mg/kg) was inhaled daily from day 1 to day 10 post infection. Mice were sacrificed at day 4 post infection and lung tissues were collected and processed to perform HE staining. Remained mice were sacrificed at day 18 post infection, which was the end point of the infection model. Lung tissues were collected, photographed and processed to perform HE staining. Body weight changes of mice was monitored through the entire infection period. (B) Photograph of lung tissues collected at day 18 post infection. Black arrows indicated obvious hemorrhagic alterations. (C) HE staining of lung tissues collected at day 4 post infection. Black arrows indicated obvious hemorrhagic alterations. Black asterisks indicated immune cells infiltration. (D) HE staining of lung tissues collected at day 18 post infection. See also in Figure S2.

### MHV-A59 infection induced potent *Acsl1* expression

We next intended to investigate how MHV-A59 infection induced ferroptosis. Although MHV-A59 infection was reported to induce enrichment of serum iron level and ROS abundance elevation in infected mice, the executor of this process remained unclear, especially in terms of ferroptosis. We infected murine BMDMs and performed bulk RNA-seq to figure out the expression profile changes brought by MHV-A59 in innate immune cells. MHV-A59 infection led to canonical inflammatory genes upregulation, characterized by the elevated expression of genes responsible for inflammatory responses (Figure 4A). Interestingly, compared with other members of acyl-CoA synthetase long-chain family, we noticed that expression of the newly identified ferroptosis executor ACSL1 was potently induced by MHV-A59 (Figure 4B and 4C). The upregulation of *Acsl1* was further confirmed in the RNA-seq data of murine PMs infected with MHV-A59 (Figure 4D). We also noticed the elevated expression of ferroptosis biomarker *Ptgs2* after MHV-A59 infection (49) (Figure 4E). We further checked the expression of *Acsl1* in several cell lines, turning out that apparent upregulation of *Acsl1* only occurred in primary macrophages (Figure 4F). In line with this, we failed to observe protective effects of ferroptosis inhibition after MHV-A59 infection in RAW 264.7 cells, compared with that observed in primary macrophages, suggesting that these protective effects may only exist in primary macrophages (Figure S3). We thus reckoned that MHV-A59 induced ferroptosis, which was dependent of the function of ACSL1. *Acsl1* was considered as an indicator of severe sepsis in addition to *Acsl4* (50). Because TLR4 activation by ligands such as LPS was one of the dominant causes for severe sepsis in bacterial infection and TLR4 inhibition by TAK-242 alleviated fatal infection by MHV-A59 (51, 52, 4), we reasoned that *Acsl1* upregulation may be the consequence of TLR4 signaling activation. Results showed that TAK-242 treatment had no inhibitory effect on MHV infection enhanced *Acsl1* expression (Figure 4G). Considering that NF-κ B activation was responsible for the explosive expression of various inflammatory genes and NF-κB inhibition suppressed *Acsl1* expression in hepatocellular carcinoma cells (53), we checked whether NF-κB inhibition attenuated *Acsl1* expression. As a result, the NF-κB inhibitor JSH-23 suppressed MHV infection induced *Acsl1* upregulation (Figure 4H). Together, these data suggested that MHV-A59 infection led to potent *Acsl1* upregulation in murine macrophages.

**Figure 4.**
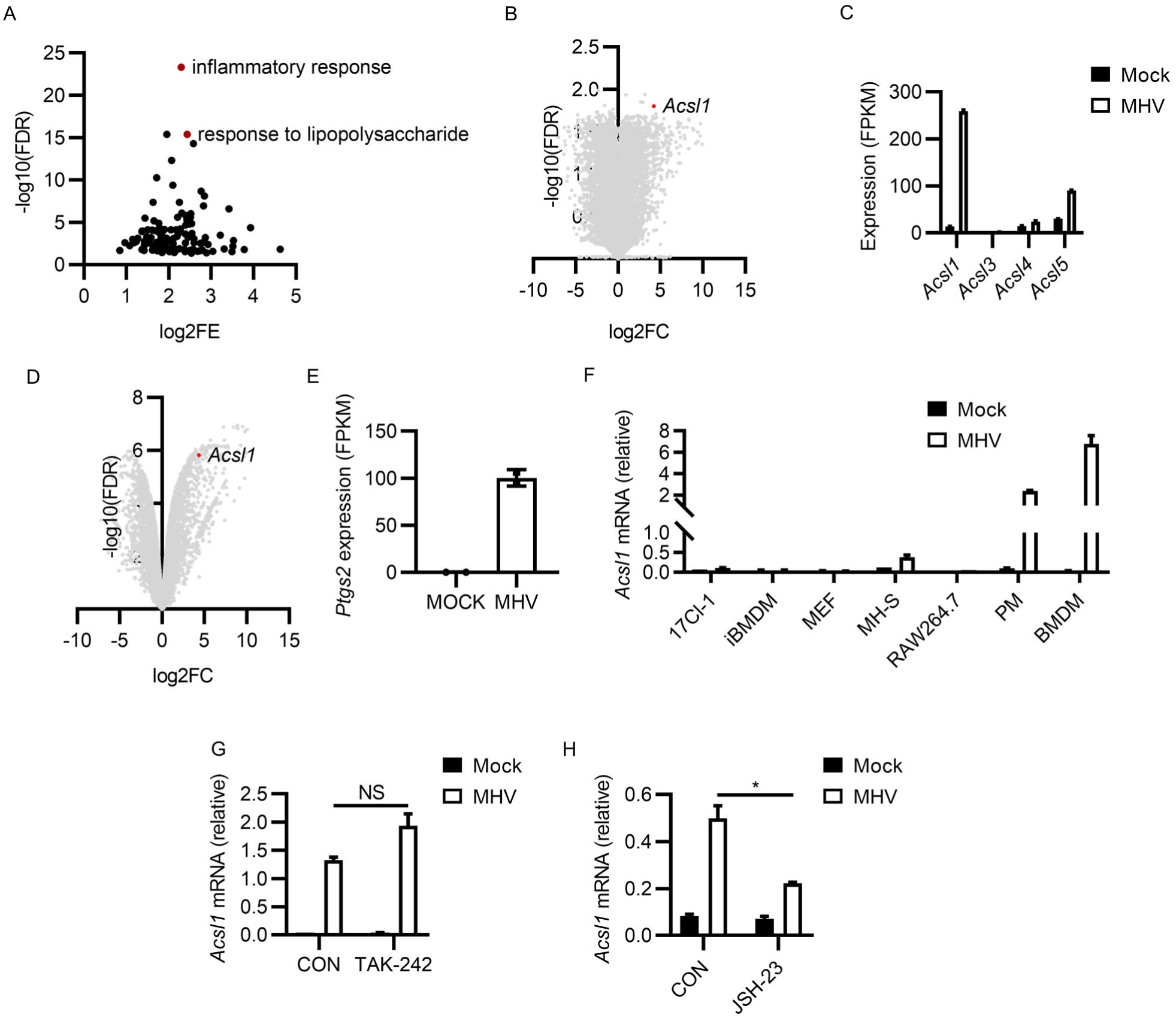
MHV-A59 infection upregulated NF-κB dependent *Acsl1* expression. (A and B) BMDMs were infected by MHV-A59 at 0.1 MOI for 12 hours and RNA was isolated to perform bulk RNA-seq. Top upregulated genes were uploaded to DAVID Bioinformatics Resources 6.8 to perform Gene Ontology analysis. Top enriched items (*p* < 0.05) were shown in (A). Expression changes of all genes (MHV infected versus control cells) was shown in (B), ranked by log2 fold change (log2FC) and –log10(false discovery rate) (-log10(FDR)). Dot of *Acsl1* was marked with red. (C) Expression of acyl-CoA synthetase long-chain family members from (A) was compared via qRT-PCR. (D) Expression changes of all genes from BMDMs after MHV infection was shown. Dot of *Acsl1* was marked with red. (E) Expression of *Ptgs2* in (D) was shown. (F) Expression of *Acsl1* after MHV-A59 infection in various cell lines was tested. (G and H) Impacts of TAK-242 (1 μM) (G) or JSH-23 (20 μM) (H) on the expression of *Acsl1* after MHV-A59 infection in PMs. Data from two independent experiments was shown. *, *p* < 0.05; **, *p* < 0.01; Student’s *t-*test.

### Inhibiting acyl-CoA synthetase long-chain family members suppressed MHV-A59 infection

Base on the results above, we suspected that targeting ACSL1 may help eliminate MHV-A59 infection. Two individual compound screening studies have found that Triacsin C inhibited SARS-CoV-2 replication (54, 55), supporting our hypothesis. We first determined the effects of ACSL1 inhibitor Triacsin C on MHV-A59 infection in murine macrophages. Triacsin C treatment significantly suppressed MHV-A59 propagation in both PMs and BMDMs (Figure 5A and 5B). In addition, Triacsin C also exerted little impact on inflammatory cytokines expression after MHV infection, similar with ferroptosis inhibitors (Figure 5C and 5D). Furthermore, Triacsin C reduced MHV-A59 induced syncytia formation (Figure 5E and 5F, see also Figure S4). These results suggested that ACSL1 inhibitor Triacsin C suppressed MHV-A59 infection.

**Figure 5.**
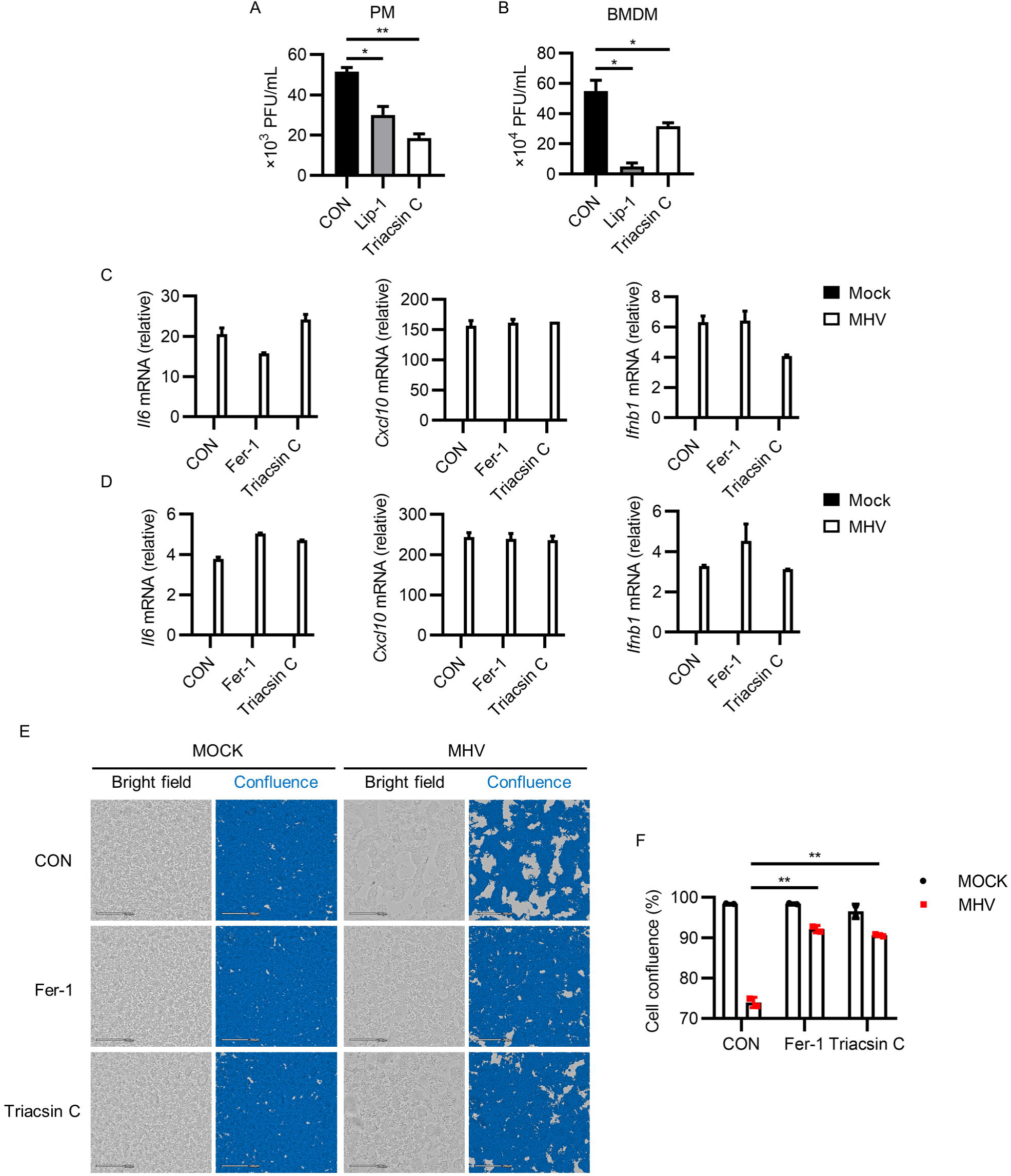
Triacsin C inhibited MHV propagation and infection induced syncytia formation. (A and B) Viral load of MHV-A59 from supernatants of PMs (A) or BMDMs (B) after Lip-1 (10 μM) or Triacsin C (2 μM) treatment. (C and D) Expression of *Il6, Cxcl10* and *Ifnb1* after MHV infection in PMs (C) or BMDMs (D) after Fer-1 (10 μM) or Triacsin C (2 μM) treatment was tested by qRT-PCR. (E and F) PMs were infected by MHV-A59 followed by treatment with Fer-1 (10 μM) or Triacsin C (2 μM). After 24 hours of infection, PMs were imaged using CytoSMART system (E) and cell confluence was evaluated using the CytoSMART website (F). Scale bars: 200 μm. Data from two independent experiments was shown. *, *p* < 0.05; **, *p* < 0.01; Student’s *t-*test. See also Figure S3.

## Discussion

In this study, we found that ferroptosis inhibitors alleviated MHV-A59 infection induced inflammation, cell syncytia formation and prevented cell death. MHV-A59 triggered potent *Acsl1* expression in murine macrophages and acyl-CoA synthesis inhibitor Triacsin C efficiently suppressed MHV-A59 infection. Our findings provided evidence for targeting ferroptosis inhibition in treating coronavirus infection related diseases.

Ferroptosis has been linked with inflammatory responses. Ferroptosis inhibition was reported to protect hepatocytes from necrotic death and suppressed immune cells infiltration in mouse nonalcoholic steatohepatitis (NASH) (57). The pro-inflammatory ligand LPS induced lipid peroxidation and injury of myofibroblast, which were inhibited by Fer-1 (58). The pro-inflammatory effects of ferroptosis can be attributed to the functional alterations of cells undergoing ferroptosis. On one hand, for immune cells, ferroptosis usually functioned via regulating their traditional activity. For example, ferroptosis in T cells suppressed T cell expansion and prevented viral clearance (59). Ferroptosis was involved in Mycobacterium tuberculosis induced necrotic cell death of macrophages and was inhibited by Fer-1 (60). On the other hand, ferroptosis in somatic cells can trigger immune cell infiltration, leading to inflammation. One example is that ferroptotic cells release HMGB1, a kind of damage-associated molecular pattern molecules (DAMPs), and mediates inflammation in macrophages (29). Given the facts above, we speculated that ferroptosis inhibition may contribute to inflammation suppression in excessive inflammation, especially in terms of coronavirus infection. SARS-CoV-2 infection can result in systemic hyper-inflammation in severe COVID-19 patients, which may be partially due to occurrence of ferroptosis. Several reviews mentioned about the possibility of ROS elimination for treatment of COVID-19. Although in one case, the application of N-acetylcysteine (NAC), a ferroptosis inhibitor that promotes glutathione supplementation, restored glutathione levels in the SARS-CoV-2 infected cells and markedly reduced C-reactive protein (CRP) level in a severe COVID-19 patient, another randomized clinical trial indicated that high dose of NAC brought little benefit to retarding the evolution of severe COVID-19 (61, 62, 63). However, our results here showed that ferroptosis inhibitor suppressed inflammatory cytokines release and MHV-A59 propagation in murine macrophages. In addition, we found that ferroptosis inhibition protected mice from MHV caused lung injury *in vivo*. Our findings broadened the application of ferroptosis inhibitors in treatments of inflammatory diseases, especially the hyper-inflammation status in coronavirus infection.

Ferroptosis is a kind of necrotic cell death characterized by the damages on cell membrane. Cell membrane rupture in lytic cell death can lead to the release of cellular components independent of canonical secretory systems, including LDH, which has been reckoned as a marker for evaluating the cell viability in lytic cell death, and cytokines such as IL-1B, TNF-α, IL-6 and CXCL-10. Differed from necroptosis and pyroptosis, until now, cell membrane rupture in ferroptosis is considered as the result of a chemical process named phospholipid peroxidation, compared with MLKL-dependent or Gasdermins-dependent pore formation. Phospholipid peroxidation on cell membrane can be reversed by free radical elimination or radical-trapping by compounds such as Fer-1 and Lip-1. Our results showed that Fer-1 and Lip-1 addition reduced viral load of MHV in cell culture supernatant, whereas the cellular level of MHV RNA was not significantly affected. This may be attributed to the reduced cell membrane damage, which may help prevent the release of viral particles. However, former studies claimed that MHV release from infected cells was mainly through lysosomes, raising the importance of inhibiting lysosome-dependent egress path. We propose that inhibiting both ferroptosis and lysosome trafficking can provide great assistance on restricting coronavirus invasion and propagation. Besides, reduced inflammatory cytokines release owing to the protective effects of ferroptosis inhibitors on cell membrane can contribute to the limitation of inflammation. Taken together, these effects indicated that cell membrane protection brought by ferroptosis inhibition can be considered as a method to restrict coronavirus infection.

ACSL1 plays important roles in regulating fatty acid metabolism in adipose tissue, liver, heart and many other tissues and organs. In immune system, upregulated *Acsl1* expression promoted inflammatory phenotype of macrophages isolated from mouse models of type 1 diabetes (64). It is possible that high level of *Acsl1* expression induced by coronavirus infection can directly enhance inflammatory phenotypes of infected macrophages via remodeling of lipid metabolism. Although inhibiting fatty acid synthesis was reported to block SARS-CoV-2 replication (65), it is still unclear how coronavirus infection alters the cellular lipid metabolism, especially the involvement of acyl-CoA synthetase long-chain family members. As a member of acyl-CoA synthetase long-chain family member,ACSL1 was recently identified to promote α-eleostearic acid triggered ferroptosis. However, the relationship between ferroptosis and inflammatory phenotypes in coronavirus infected macrophages is still elusive. Apart from macrophages, ACSL1 was also observed to be highly expressed in neutrophils in fatal sepsis (50). Our *in vivo* data revealed that ferroptosis inhibition protected mice from MHV infection,. The protective effects may be ACSL1-dependent. Taking the sepsis-like symptoms of COVID-19 patients into consideration, ACSL1 may serve as an intriguing therapeutic target to restrict cytokine storm in COVID-19.

Upregulation of *Acsl1* was reported to depended on Toll-like receptors and NF-κB. In our experiments, we evaluated the effects of TLR4 inhibitor TAK-242 and NF-κB inhibitor JSH-23 on expression of *Acsl1*. We observed that induction of *Acsl1* in MHV infected macrophages was NF-κB-dependent and TLR4-signaling-independent. Although our previous study indicated the importance of TLR4 signaling in restricting MHV infection, our results here showed that in macrophages, *Acsl1* upregulation required NF-κB but not TLR4 activation. Because Acsl1 expression also depends on TLR2 and TLR2 deficiency renders mice more protected from MHV infection (56, 66), we speculate that TLR2 signaling is responsible for MHV induced *Acsl1* expression and plays more important roles than TLR4 signaling in mediating MHV infection. Different from SARS-CoV-2 infection, which was reported to downregulate *GPX4* expression, however, MHV infection resulted in no significant reduction of *Gpx4* expression, indicating distinct routines of ferroptosis occurrence between SARS-CoV-2 and MHV infection.

Triacsin C is a pan-acyl-CoA synthetase inhibitor which exerts inhibitory effects on ACSL1, ACSL3 and ACSL4. For macrophages infected by MHV, we showed that ACSL1 constituted the dominant isoform of acyl-CoA synthetase family members. Therefore, Triacsin C was considered as the ACSL1-sepcific inhibitor in our experiments. Two independent compound screening studies showed that Triacsin C can be a candidate for restricting SARS-CoV-2 infection, further validating the involvement of acyl-CoA synthetase family members in coronavirus infection. However, due to the lacking of target specificity, it is challenging for Triacsin C to be applied for the inhibition of ACSL1 *in vivo*. Recently, a compound based on Triacsin C showed highly potent and selective inhibitory effects on ACSL1 (68). This refined compound may be selected as an ideal inhibitor for treatment of coronavirus infection.

Collectively, our study provided the implications for focusing on the correlation between ferroptosis and inflammation in coronavirus infection. Our findings emphasized the importance of acyl-CoA synthetase family members, especially ACSL1 in MHV infection, and showed that ferroptosis inhibition offered benefits for individuals suffering coronavirus infection.

## Materials and methods

### Cells

RAW 264.7, iBMDM, MEF and 17CL-1 cells were kept in our lab. Mouse alveolar macrophage cell line MH-S cells were from American Type Culture Collection (ATCC). Cells were cultured in Dulbecco’s modified eagle medium (DMEM) supplemented with 10% FBS (PAN), 1% glutamine (Gibco), 100 U/ml penicillin and 100 μg/ml streptomycin at 37°C in the incubator ESCO^®^ CCL-170B-8 with the presence of 5% CO_2_.

### Primary macrophages

For the isolation of mouse primary peritoneal macrophages (PMs), cells were collected from lavage of the peritoneal cavity from mice that were pre-stimulated with thioglycolate (TG) for 3 days. For the isolation of bone marrow derived macrophages (BMDMs), bone marrow was rinsed out of femurs and tibiae of 6-week-old C57BL6/ mice. Cells were cultured with 30 % L929 cell culture supernatant for 3 days. 3 days later, the supernatant was replaced by fresh cell culture medium with 30% L929 cell culture supernatant and maintained for another 3 days. Mature macrophages were harvested by digesting with trypsin-EDTA for 2 minutes and seeded into 24-well plates for further experiments (200 thousand cells per well).

### Murine hepatitis virus culture

The murine hepatitis virus A59 strain has been described previously (2). MHV-A59 was propagated in 17CL-1 cells. Briefly, 0.005 multiplicity of infection (MOI) MHV-A59 was added to the supernatant of growing cells. The cell debris and supernatants were collected, resuspended and froze in −80°C freezer 36 hours later. The supernatant was frozen and thawed for 3 cycles, centrifuged and subpackaged, followed by the assessment of virus titer via plaque assay.

### Reagents and drugs

Ferrostatin-1 (HY-100579) and TAK-242 (HY-11109) were from MedChemExpress. Liproxstatin-1 (S81156) was from Yuanye Bio-Tech. Triacsin C (T139793) was from Aladdin. VX-765 (T6090), z-DEVD-FMK (T6005), GSK-872 (T4074), and JSH-23 (T1930) were from Targetmol.

### Virus infection in cells

RAW264.7 cells, PMs and BMDMs were infected by MHV-A59 at indicated MOI. 2 hours after infection, the supernatant was removed and cells were covered with culture medium containing compounds at indicated concentration. Infected cells or supernatants were harvested to perform the further experiments.

### RNA isolation and RT-qPCR

Total RNA from infected cells was extracted via one-step method using Trizol reagent (Invitrogen) and the cDNA was generated using HiScript II Q RT SuperMix (Vazyme). Quantitative real-time PCR was carried out using ChamQ Universal SYBR qPCR Master Mix (Vazyme) and specific primers on the 7500 Fast Real-Time PCR Instrument (Applied Biosystems). Data were analyzed using GraphPad Prism Version 8.0 (GraphPad Software) according to the 2^-ΔCt^ threshold calculation method and means ± SD. The relative RNA expression level was normalized to *Hprt* quantified in parallel amplification reactions. Primer sequences were listed as follows:

m*Hprt*-F: 5’-TCAGTCAACGGGGGACATAAA-3’

m*Hprt*-R: 5’-GGGGCTGTACTGCTTAACCAG-3’

m*Il6*-F: 5’-TCTGCAAGAGACTTCCATCCAGTTGC-3’

m*Il6*-R: 5’-AGCCTCCGACTTGTGAAGTGGT-3’

m*Ifnb1*-F: 5’-TCCGAGCAGAGATCTTCAGGAA-3’

m*Ifnb1*-R: 5’-GCAACCACCACTCATTCTGAG-3’

m*Cxcl10*-F: 5’-CCAAGTGCTGCCGTCATTTTC-3’

m*Cxcl10*-R: 5’-GGCTCGCAGGGATGATTTCAA-3’

m*Acsl1*-F: 5’-TGCCAGAGCTGATTGACATTC-3’

m*Acsl1*-R: 5’-GGCATACCAGAAGGTGGTGAG-3’

MHV-pp-F: 5’-TGCCTGAAACGCATGTTGTG-3’

MHV-pp-R: 5’-CAGACAAACCAGTGTTGGCG-3’

### Bulk RNA-seq

Integrity of RNA purified from indicated BMDM or PM samples was assessed using the RNA Nano 6000 Assay Kit of the Agilent Bioanalyzer 2100 system (Agilent Technologies). All samples showed RNA integrity number > 8. RNA sequencing libraries were generated using NEBNext Ultra Directional RNA Library Prep Kit for Illumina (NEB), and sequenced on an Illumina Novaseq PE150 platform. Sequencing was performed at Genewiz Co. Ltd. The filtered reads were mapped to the mouse genome reference sequence (GRCm38/mm10 Ensembl release 81) using HISAT2. Gene expression was quantified as fragments per kilobase of coding sequence per million reads (FPKM) algorithm. Genes were ranked by log2 of fold change (log2FC) and -log10 of false discovery rate (-log10(FDR)). For the gene ontology analysis, top upregulated genes were uploaded to DAVID Bioinformatics Resources 6.8 to perform Gene Ontology (GO) analysis. Top enriched items (*p* < 0.05) were shown in the figure, as described in the legend of figure 4.

### PI staining

Real-time cell membrane permeability assessment was carried out using PI staining. Live cells were covered by cell culture medium with 20 μM PI. Cells were cultured for 15 min followed by washing with PBS for 2 times. Then cells were covered with culture medium and monitored under fluorescence microscope.

### Cell imaging

For the imaging under bright field alone, cells were monitored using CytoSMART Lux2 cell imaging system (CytoSMART). Extent of syncytia formation was determined with cell confluence level, which was calculated in CytoSMART website. For the imaging of cells stained with PI, cells were monitored using Olympus IX70 Fluorescence Microscope (Olympus).

### Cytokine measurement and LDH assay

For the detection of cytokine abundance in the cell culture supernatant, supernatants from treated cells were collected to perform ELISA following the manufacturer’s instructions. The following ELISA kits was used: mouse IL-6 ELISA kit (Dakewei). LDH was measured via colorimetric NAD linked assay using LDH detection kit (Leagene).

### Plaque assay

Virus yield of MHV-A59 in culture supernatants was determined by plaque assay with 17CL-1 cells. Culture supernatant was harvested and diluted to infect confluent 17CL-1 cells cultured in 24-well plates. Viruses were removed 2 hours after infection and cells were washed with pre-warmed PBS followed by culture with DMEM containing 0.5% methylcellulose. After 36 hours of infection, the overlay was removed and cells were fixed with 4% paraformaldehyde for 10 min and stained with 1% crystal violet for 20 min. Plaques were counted, averaged, and multiplied by the dilution factor to determine the viral titer as plaque-forming units per mL (PFU/mL).

### Animal model

All mice were bred and maintained in specific pathogen-free (SPF) conditions with approval by the Peking University animal care and use committee. For the infection experiments, 4-week-old C57BL/6 male mice were anesthetized via inhaling isoflurane and then inoculated intranasally (i.n.) with 10 μL of MHV-A59 virus at 1×10^4^ PFU. Mice were intranasally treated daily with Lip-1 or vehicle alone from day 1 to day 10 post infection or mock infection. We monitored the weight of each mouse every day until mice were sacrificed. Mice were sacrificed at day 4 or day 18 post infection or mock infection to evaluate the tissue injury and inflammation caused by MHV-A59 infection.

### Histology

Mice were sacrificed at indicated time points after infection. Lungs from each group were fixed with 4% paraformaldehyde, embedded in paraffin and cut into sections of 3.5 μm, and further stained with hematoxylin and eosin (HE). Immune cells infiltration and hemorrhages were determined and evaluated under light microscopy. The images were taken by Leica DM 6B microscope.

### Statistical analysis

Results were analyzed by paired or unpaired Student’s *t-*test or by two-way ANOVA analysis (for determining the differences in the weight loss curves). GraphPad Prism 8.0 was used for statistical analysis and graphing. Data was shown as mean ± SD unless indicated in the legend. Statistical values can be found in the figure legends. \**p* < 0.05, \*\**p* < 0.01, \*\*\*\**p* < 0.0001. For experiments *in vivo, n*= number of individual animals.

## Figure legends

Figure S1. Fer-1 does not inhibit viral entry of MHV-A59.

Referring to Figure 2. PMs were pre-treated with Fer-1 (10 μM) for 2 hours and infected with MHV-A59 at 1 MOI for 2 hours. Expression of MHV-pp was tested by qRT-PCR. NS, not significant.

Figure S2. Lip-1 was not able to reverse reduced gain of body weights in low-dose MHV-A59 infection model.

Referring to Figure 3. Body weight changes were monitored daily. Apparent restriction of gain of body weight compared with non-infected group was observed after MHV-A59 infection. Data was shown as means ± SEM. **, *p* < 0.01; Student’s *t-*test. NS, not significant.

Figure S3. RAW 264.7 cells were not protected from ferroptosis inhibition after MHV-A59 infection.

(A) RAW 264.7 cells were infected with MHV-A59 at 0.05 MOI for 24 hours, stained with PI and imaged under fluorescence microscope. RAW 264.7 cells showed no obvious membrane permeability alterations after MHV-A59 infection.

(B and C) RAW 264.7 cells were infected with MHV-A59 at 0.05 MOI for 24 hours. Expression of *Ifnb1* (B) and MHV-pp (C) was tested by qRT-PCR.

(D) Viral load of MHV-A59 from supernatants of RAW 264.7 cells with or without Fer-1 (10 μM) treatment. NS, not significant.

Figure S4. Triacsin C inhibited MHV induced syncytia formation in BMDM.

Referring to Figure 5. Impacts of Fer-1 (10 μM) or Triacsin C (2 μM) on syncytia formation of BMDMs after MHV-A59 infection. Black arrows indicated cell syncytia.

Video S1. MHV-A59 infection induced ferroptosis-like morphological changes of murine peritoneal macrophages

Referring to Figure 1. PMs were infected with MHV-A59 at 0.1 MOI. Cell morphology was monitored under CytoSMART Live Cell Imaging System for 24 hours.

## Acknowledgments

We thank Dr. Zhao Yingchi for help refining the viral propagation experiments. This work was supported by the National Key Research and Development Program of China (2016YFA0500300; 2020YFA0707800), the National Natural Science Foundation of China (31570891; 31872736; 32022028; 81991505), Peking University Clinical+X (PKU2020LCXQ009), the Peking University Medicine Fund (PKU2020LCXQ009.

## Author contributions

F.Y. and Z.Z. designed the research; X.H. and Z.Z. performed the experiments and analyzed data; F.Y. supervised the study; Z.Z., X.H. and F.Y. interpreted data and wrote the manuscript. All authors commented on the manuscript.

## Data and materials availability

The raw data of RNA sequencing in this publication has been deposited in NCBI’s Gene Expression Omnibus and are accessible through GEO Series accession numbers GSE185800.

## Abbreviations

MHV-A59: Murine hepatitis virus strain A59
DAMP: damage-associated molecular pattern molecule
COVID-19: coronavirus disease 2019
ROS: reactive oxygen species
NDV: new castle virus
BMDM: bone marrow derived macrophage
PMs: peritoneal macrophages
Fer-1: ferrostatin-1
Lip-1: liproxstatin-1

