## Supplementary figures and images for "Inhibiting ACSL1 related ferroptosis restrains MHV-A59 infection"

### Supplemental figure 1

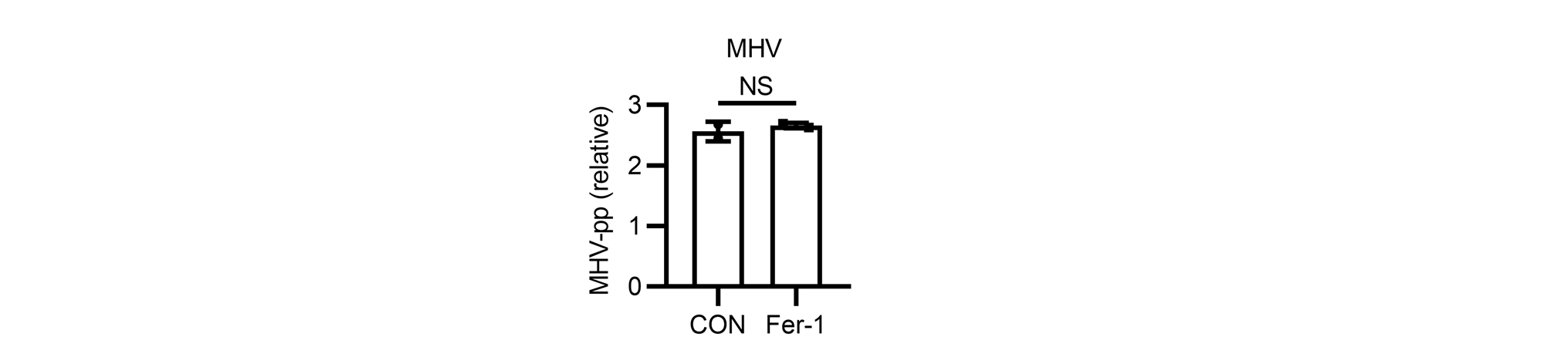

### Supplemental figure 2

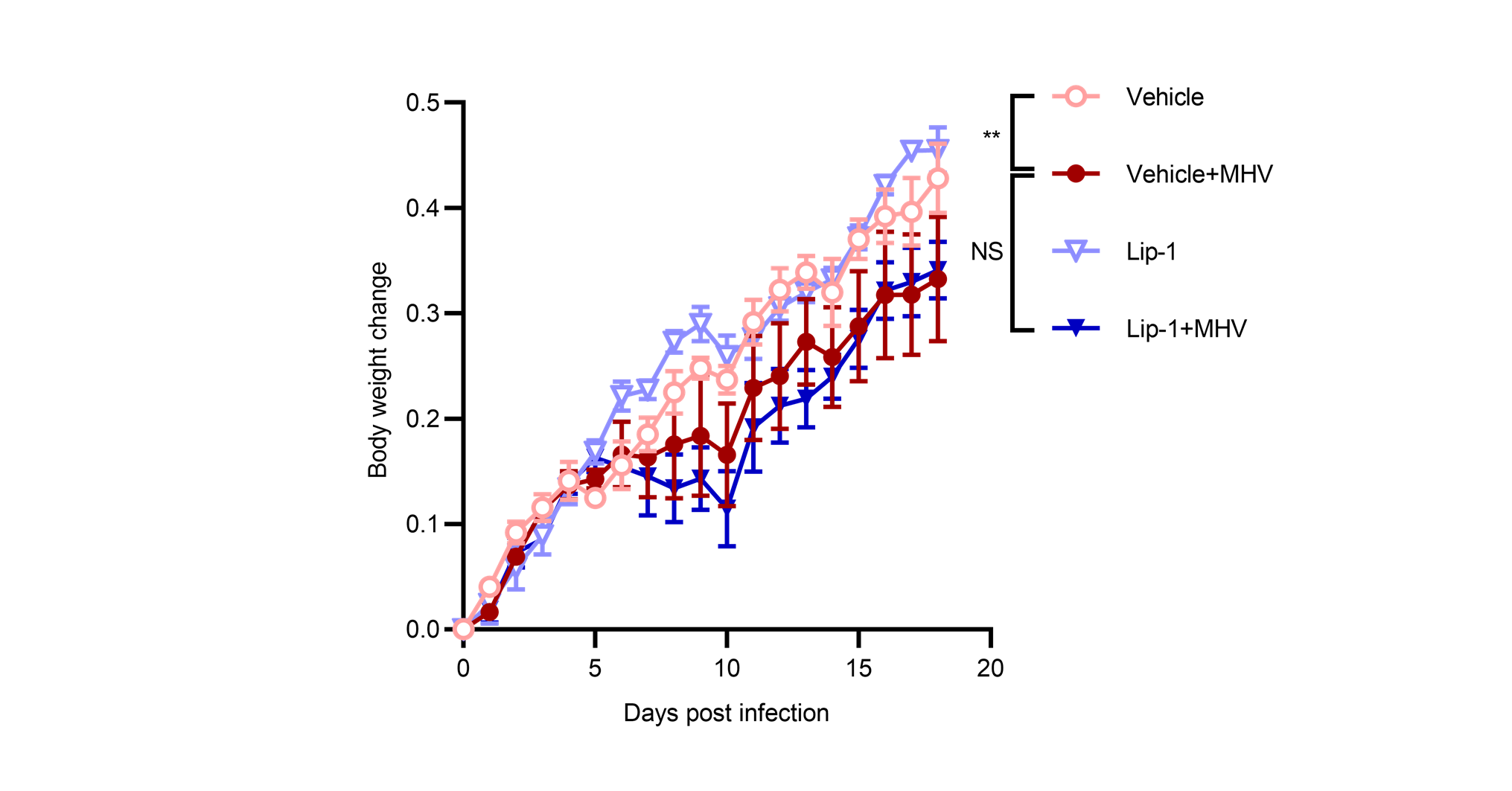

### Supplemental figure 3

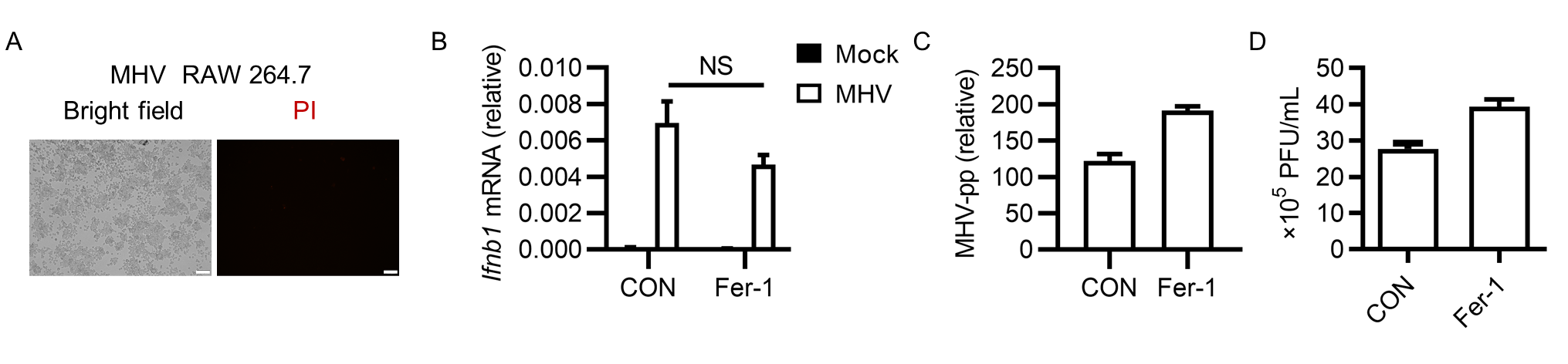

### Supplemental figure 4

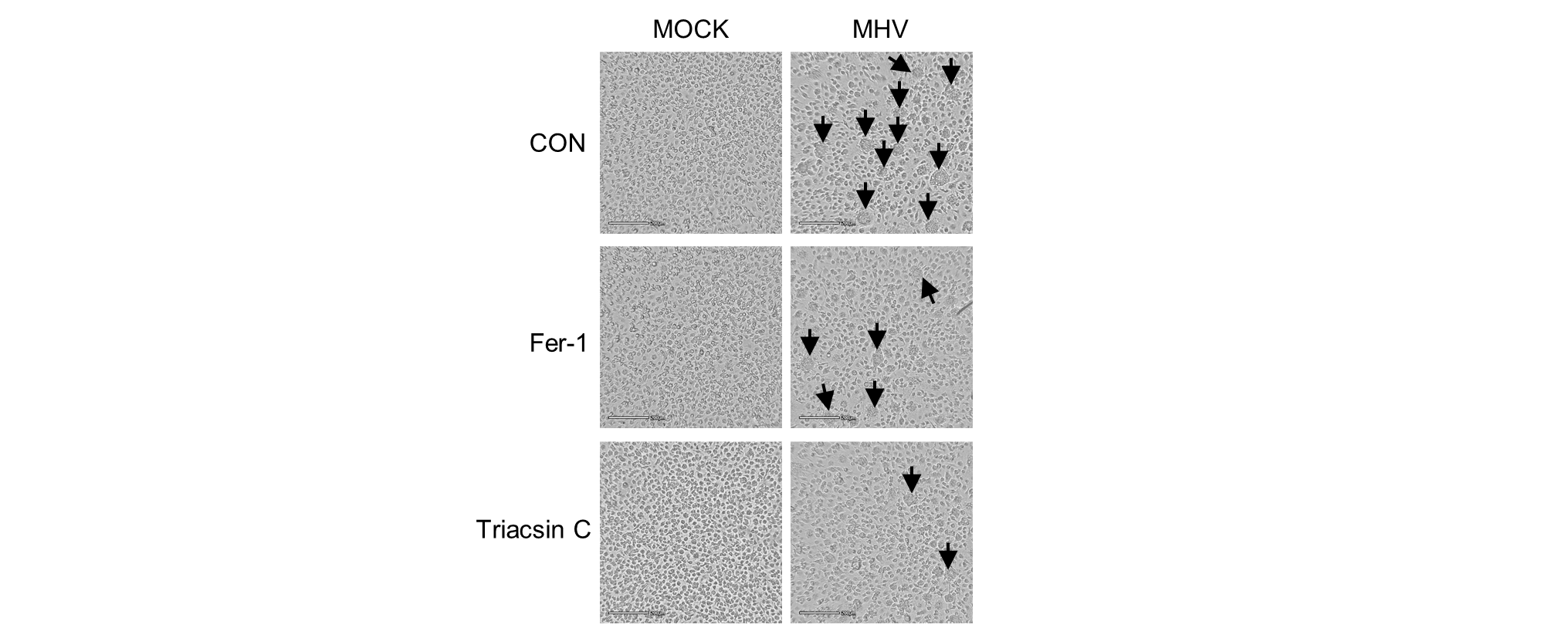
